# Kinetic ^13^CO_2_ mapping revealed distinct light-dark metabolic transition phenotypes in *Brassica napus* seedlings under visible and UV-B light

**DOI:** 10.64898/2025.12.03.692083

**Authors:** Yogita Thakur, Maneesh Lingwan, Yogesh Pant, Shyam Kumar Masakapalli

## Abstract

Dynamic stable-isotope tracing using ^13^CO_2_ has gained significant attention in systems biology due to its potential to visualise carbon assimilation patterns. However, the plant metabolic phenotypes under visible and UV-B light, explaining the light-to-dark transition between photoperiods, remain unexplored. In this study, we investigated the dynamics of photosynthetic carbon assimilation and resource partitioning during the day-night transition under visible and UV-B light in *Brassica napus* seedlings. Kinetic ^13^CO_2_ tracing via the analysis of mass isotopomer distributions of metabolite fragments using GC-MS revealed reprogramming of the source-sink carbon dynamics. While visible light enabled the dynamic redistribution of newly fixed carbon during the light and dark photoperiod, de novo biosynthesis of shikimic acid, TCA cycle intermediates and metabolites of the glutamate-GABA metabolism was strongly favoured in dark metabolism. In contrast, a delayed and reduced photosynthetic carbon assimilation response was observed in the UV-B phenotype during the light period. Moreover, towards the late light period, de novo biosynthesis of sucrose, shikimic acid, phenylalanine, citric, succinic and malic acid was favoured, along with a greater reliance on pre-existing carbon pools for other metabolites. However, ketoglutarate, succinic acid, malic acid and GABA showed limited de novo synthesis in the dark period. Across both light regimes, amino acid pools largely remained in constant sync with the pre-existing pools during the light-dark transition. Overall, our findings demonstrate that light quality and photoperiod-driven metabolic transitions distinctly shape plant metabolic phenotypes.

## Introduction

Due to their sessile nature, plants are constantly exposed to various stress factors, and thus they have to rely on their metabolic flexibility to adjust and adapt towards these changes (Kiss, 2006; Singh et al., 2021; Lingwan, 2024; Seth & Sebastian, 2024). Plants use carbon as the main currency to support their growth, defence and survival via photosynthesis. As plants experience stressful environmental conditions, there is a change in the dynamicity of carbon in the form of source-sink metabolites (Foyer & Paul, 2001; Fester et al., 2013; Bénard et al., 2015; White et al., 2016; Fernie et al., 2020; Chen et al., 2021; Rosado-Souza et al., 2023; Gessler & Zweifel, 2024; Li et al., 2025). Various environmental stresses, including the spectral light quality, can alter these carbon allocation patterns. Therefore, understanding the photosynthetic carbon assimilation and distribution patterns becomes important in defining the plant metabolic phenotypes.

Light serves as a primary energy source for photosynthesis and functions as a vital environmental signal, significantly influencing plant development and physiology (Lingwan et al., 2023; W. Wu et al., 2024; Yadav et al., 2020). The metabolic phenotype of a plant is strongly influenced by the spectral quality of light, influencing the photosynthetic carbon assimilation and partition. Due to the absorption of light wavelength by the photosynthetic pigments, plants can photosynthesise in the visible spectrum of light (400-700 nm) (Van Grondelle & Boeker, 2017; Liu & van Iersel, 2021; B. Sen Wu et al., 2024). In contrast, among its varied spectral components, ultraviolet (UV) radiation (200 – 400 nm), although comprising only about 7% of the total solar radiation, plays a significant role in modulating plant responses (Meyer et al., 2021; Job et al., 2022; Lingwan et al., 2024). In recent decades, surface UV-B radiation has shown changes regardless of variations in stratospheric ozone levels, largely due to climate-driven shifts (Williamson et al., 2014; Barnes et al., 2019; Bernhard et al., 2023). The effects of UV-B on plants are complex, ranging from beneficial to detrimental (Jansen et al., 1998; Frohnmeyer & Staiger, 2003; Hideg et al., 2013; Z. S. Zhang et al., 2016; Fraser et al., 2017; Qi et al., 2018; Yadav et al., 2019; Saini et al., 2020; Shi & Liu, 2021; J. Sun et al., 2023; Job N. et al., 2025). Therefore, plant responses to UV-B are highly variable, often involving an extensive metabolic reprogramming within the plant system to maintain a state of homeostasis.

Besides understanding the highly variable metabolic responses of plants to the spectral light quality, the day-night transition presents an additional layer of complexity. Several studies have shown the utilisation of different carbon pools during the light and dark phases of the photoperiod and its regulation via the circadian rhythms (Lawlor & Fock, 1977; Dohler, 1989; Tronconi et al., 2008; Graf et al., 2010; Zell et al., 2010; Graf & Smith, 2011; Tcherkez et al., 2012; Szecowka et al., 2013; Müller et al., 2014; Sulpice et al., 2014; Kölling et al., 2015; MacNeill et al., 2017; Abadie & Tcherkez, 2019; Abadie et al., 2024).

While a static overview of plant metabolism can be obtained with metabolomics (Meiser & Frezza, 2024), investigating the dynamic flow of carbon in plant metabolism can be aided by stable isotope labelling studies. The choice of isotopic tracer depends on the biological question of interest (Koley et al., 2024). ^13^CO_2_ has been demonstrated as a tracer for studying autotrophic tissue metabolism, while ^13^C sugars and amino acids can be employed to trace the flow of carbon within a mixotrophic tissue. For exploring the metabolism of heterotrophic tissue, ^15^NO_3_ and ^2^H_2_O are widely used.

Belonging to the *Brassicaceae* family, rapeseed is a major source of edible oils (FU et al., 2016; Tassone et al., 2016; Begna et al., 2017; Weselake et al., 2024; Mohamed et al., 2025). Approximately 11% of the global rapeseed-mustard (*Brassica napus*) production comes from India, making it the third-largest producer of this oilseed crop (Chongloi et al., 2025), highlighting the importance of rapeseed in ensuring global food security. Rapeseed is also the second most cultivated oilseed crop in India, after soybeans, and in terms of oil production, it ranks second internationally (Borges et al., 2023). Rapeseed oil is enriched with many bioactive compounds beneficial for human health (G. Yang et al., 2025; J. M. Yang et al., 2024), and even the by-products of rapeseed oil production, including rapeseed cake, act as excellent nitrogen-rich organic fertilisers (Liang et al., 2025). *Brassicaceae* microgreens have also been established as a promising functional food (Pant et al., 2023).

One of the most critical stages in a plant’s life cycle is the seedling stage (Hanley et al., 2004), as its survival greatly determines a plant’s growth. Therefore, this stage plays an important role in shaping the morphological and physiological aspects of plant populations. It is the stage when the plant initiates active photosynthesis, leading to an independent life cycle (Cao et al., 2022). Therefore, it becomes crucial to understand the changes occurring within the plant system during this particular phase of development. Hence, this stage becomes crucial for plant survival and agricultural production (Ali et al., 2021). Thus, understanding the metabolic changes during this window becomes important in gaining insights into the early metabolic adjustments, as this stage is very sensitive to environmental changes.

Most studies have explored the morphological, biochemical, and gene expression level changes occurring in *Brassica* sp. on UV exposure (Lee et al., 2022; Gray et al., 2024); no studies have investigated the metabolic adaptations to UV-B at the seedling stage using stable isotopic mapping in *Brassica napus* (Gray et al., 2024; Lee et al., 2022). Jing et al. (2023) described the transcriptional and metabolomic changes in 2 and 8-week-old *Brassica* varieties exposed to UV-B. (Dellero et al., 2023) in their work focused on defining the metabolic fluxes in *Brassica napus* leaves under normal growth conditions with 6 hours of labelling. Schwender & Ohlrogge (2002) emphasised the storage metabolism of *Brassica* embryos using a stable isotope labelling approach. However, comprehensive studies integrating prolonged ^13^CO_2_ mapping of metabolites and their labelling dynamics under visible and UV-B light explaining the transition between photoperiods remain unexplored. Therefore, this study aims to utilise kinetic ^13^CO_2_ mapping to define the metabolic phenotype and light-to-dark metabolic transitions in *Brassica napus* var. *sheetal* seedlings under visible and UV-B light conditions, to understand the carbon allocation and partitioning criteria. Our findings will also highlight the behaviour of stored (unlabelled) vs. newly fixed (labelled) carbon during the extended light-to-dark phase transition under visible and UV-B light.

## Materials and Methods

### Plant material and growth conditions

Seeds of *Brassica napus* var. *sheetal* were acquired from CSK Himachal Pradesh Agricultural University, Palampur. Surface sterilisation of seeds for two minutes was done using 1 % sodium hypochlorite solution. A thorough washing of surface-sterilised seeds using distilled water was done five times to remove any traces of mercuric chloride. The seeds were germinated in an autoclaved cocopeat-vermiculite substrate mixture (1:1 v/v). For the initial three days, the seeds were allowed to germinate in the dark at an ambient temperature maintained at 22 ± 2 °C and relative humidity levels of 60 ± 5 %. On day four, seedlings were transferred to light (16:8 light/dark photoperiod) and allowed to grow till they reached the 8-day-old seedling stage. The eight-day-old seedlings were subjected to UV-B and visible light conditions for 16 hours light phase followed by 8 hours dark phase (24 h). The photosynthetic photon flux density for visible light experiments was set to 90-110 µmol m^−2^ sec^−1^ using the growth chamber Percival LED22C8, and for the UV-B experiment, a Phillips TL20W/01RS SLV narrow band UV-B lamp (305-315nm) having λ_max_ at 311 nm was used. For measuring the intensity of visible and UV-B light, Apogee Quantum meter MQ-200 and UVA/UVB–850009 (Sper scientific) light meters were used (Lingwan et al., 2024).

### Kinetic labelling of plants using ^13^CO_2_

A parallel ^13^CO_2_ feeding chamber prototype was developed and optimised in 16:8-hour photoperiodic conditions for ^13^CO_2_ feeding experiments under visible and UV-B light conditions (Lingwan et al., 2024). Using three different mass flow controllers, CO_2_/^13^CO_2_, N_2_ and O_2_ gases from different cylinders were supplied to the gas mixing system in a controlled manner for the proper mixing of gases to create synthetic air. The levels of CO_2_ were maintained at 400 ppm ± 50 ppm. To ensure the complete equilibrium of ^13^CO_2_ after a step change, the chamber dead volume (which accounts for pre-existing ^12^CO_2_) was calculated. The size and volume (33 cm (L) × 24 cm (H) × 22cm (W); ∼12.6 L) of the boxes were kept the same to avoid any experimental bias. The volumetric flow rate was 0.4998 L/min with constant internal and external temperature. To track the temperature and humidity, sensors were used, and kinetic sample quenching was done in liquid nitrogen at 0, 0.5, 1, 2, 4, 8, 16, and 24 hours. The chamber design enabled rapid quenching of metabolism without environmental exposure.

### Metabolite extraction and GC-MS profiling

Rapeseed seedling samples were quenched in liquid nitrogen, crushed and lyophilised, and ∼20mg of this lyophilised sample was used for the extraction of soluble metabolites using the extraction solvent. Extraction solvent contained a 3:1:1 ratio of methanol, chloroform and water. To 940 µl of extraction solvent, 60 µl of ribitol (0.2 mg ml^-1^ stock in water) was added per sample. The mixture was subjected to a temperature of 70°C in a thermoshaker at 950 rpm for 5 minutes, followed by centrifugation of extracts at 13000 g at room temperature for 10 minutes (Lisec et al., 2006). An aliquot of supernatant (50 µl) is transferred to a new eppendorf tube and subjected to speed vacuum drying. MSTFA (N-Methyl-N-Trimethylsilyl-trifluoroacetamide) was used for TMS (trimethylsilyl) derivatisation of the dried samples (Lingwan & Masakapalli, 2022; Masakapalli et al., 2013, 2014). For data acquisition, the samples were subjected to GC-MS using Agilent Technologies GC ALS-MS 5977B with having HP-5MS (5% phenyl methyl siloxane) column (30 m x 250 µm x 0.25 µm). The sample injection volume was 1 µL in splitless mode. 70eV electron ionisation was used. Other program measures, including temperature program and scan mass-range parameters, were set as mentioned by Lingwan & Masakapalli (2022).

### ^13^C isotopomer profiling using GC-MS

Metalign software was used for baseline correction of the raw GC-MS spectral files (Lommen, 2009; Lommen & Kools, 2012). Profiling of metabolites by identification of spectral peaks was done using Agilent ChemStation software by comparing the mass ion fragment (m/z) and retention time (RT) against the NIST 17 library (Schauer et al., 2005) and in-house standards. Metabolites having >70% probability score in the NIST 17 library were chosen for further analysis. For ^13^C incorporation in the metabolites, the intensities of the mass fragments of amino acids and organic acids of primary metabolic pathways were obtained using Agilent ChemStation software and the fragments were corrected for the natural abundance of isotopes in the carbon backbone using IsoCor software (Millard et al., 2012). The organic acid and amino acid derivative and metabolite formulas were calculated for TMS-derivatised soluble metabolites using a correction method explained in **Supplementary Table S1**. For further analysis, only the valid fragments were considered to determine the average ^13^C incorporation. Additional fragments generated during this study, with their average ^13^C incorporation, are listed in the **Supplementary Table S2**.

### Statistical analysis

For statistical analysis, significance was determined by unpaired t-tests wherever required. GraphPad Prism 10.3.1 and Microsoft Excel were used for data visualisation. Error bars were expressed as the standard error of the mean (SEM) for four biological replicates.

## Results and discussion

### Kinetic ^13^CO_2_ mapping revealed divergent light-driven metabolic phenotypes

The metabolic profiling of soluble extracts of 8-day-old rapeseed seedlings exposed to visible and UV-B light captured 48 metabolic features (**Supplementary Table S3)**. The levels of these metabolites fluctuated in visible and UV-B light conditions. Metabolites associated with central carbon metabolism and the shikimic acid pathway were selected for the kinetic ^13^CO_2_ mapping.

On exposure to visible light, key intermediates of the primary metabolic pathway, such as the tricarboxylic acid (TCA) cycle intermediates (citric acid, keto glutarate, succinic acid and malic acid) and amino acids (serine, phenylalanine, valine, leucine, glutamic acid, isoleucine, threonine, GABA, and oxoproline), exhibited a general trend of consistently increased ^13^CO_2_ incorporation (**Figure 1)**. Sucrose is observed to be the major photosynthetic product (Jensent et al., 1966; King et al., 1967; Stitt & Ap Rees, 1980; Szecowka et al., 2013) with an average ^13^C of 33.7% which is the highest label incorporation among all other detected metabolites. The kinetic data of all the examined metabolites showed an increasing trend of ^13^C label incorporation during the initial hours of photosynthesis (till 8h), after which there is a decline in label incorporation until the end of the light period (16h). This indicates that there is an active ^13^CO_2_ assimilation followed by dilution during this photoperiod. Between 0 to 8h, the percentage of label incorporation in the major photosynthetic product, sucrose, ranged from 0.6 to 32.3%. These observations suggest that under typical visible light, there is efficient carbon fixation. The metabolic network remains highly active, efficiently converting fixed carbon into sucrose, which subsequently feeds into the glycolytic intermediates.

**Figure 1:**
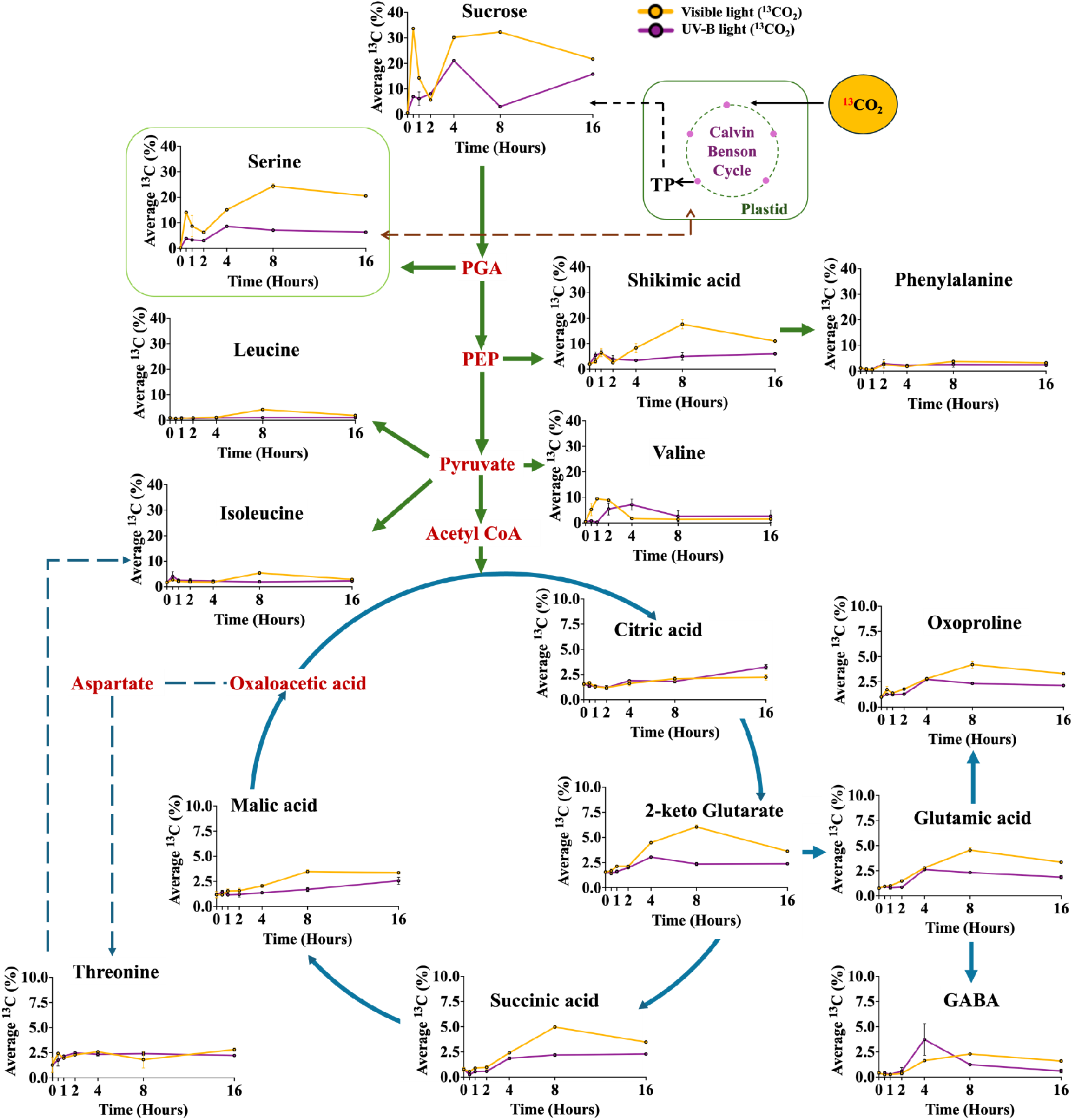
Kinetic 13C enrichments of soluble metabolites highlight metabolic readjustments. Average 13C incorporation in soluble metabolites of 8-day-old rapeseed seedlings exposed to visible light and UV-B light kinetically. The orange line represents the average ^13^CO_2_ incorporation under visible light conditions, and the violet line represents the average ^13^CO_2_ incorporation under UV-B light conditions. The average ^13^CO_2_ incorporation was lower under UV-B light compared to visible light conditions. Error bars are expressed as the standard error of the mean (SEM), where n=4. **Supplementary Figure S1** represents the labelling pattern in the central metabolic pathway intermediates, showing the average 13C incorporation (%) in TMS fragments.

The TCA cycle intermediates exhibited a gradual increase in ^13^C label incorporation till the mid-light phase (8h), followed by the label dilution during the late-light period (16h). The highest label incorporation was observed in ketoglutarate, which was about 6%. There is an efficient carbon flow from ketoglutarate to glutamic acid, which further channels the fixed carbon to oxoproline and GABA. Overall, the results indicate that under normal illumination conditions, there is an active assimilation of ^13^CO_2_ into the central metabolites via glycolysis, TCA cycle and glutamate-GABA metabolism.

In contrast, under UV-B light, the incorporation of ^13^CO_2_ in the above-mentioned metabolic intermediates is substantially reduced, and this effect is nearly uniform for almost all metabolites **(Figure 1)**, indicating that UV-B stress might affect carbon uptake and downstream metabolism (Dohler, 1989; Kaling et al., 2015). An altered pattern of label incorporation was observed in the seedlings exposed to UV-B. Some metabolites showed a trend of increased label incorporation until 4h (sucrose, serine, valine, ketoglutaric acid, glutamic acid, oxoproline and GABA) after which there is an observed trend of label reduction. Other metabolites such as shikimic acid, phenylalanine, leucine and isoleucine showed a spike in label incorporation in the much earlier stages of light exposure (1-2h). The TCA cycle intermediates, including citric acid, succinic acid and malic acid, showed a steady increase in label incorporation up to the 16h time point. This suggests that the metabolic network is unable to maintain its high level of photosynthetic activity under UV-B light, and as a result, carbon uptake and subsequent downstream metabolic processes are hampered. There may be stress-induced metabolic readjustments (Höll et al., 2019) leading to metabolic suppression. Under UV-B light, the plant may prioritise defence mechanisms over energy production (Höll et al., 2019; Shi & Liu, 2021; H. Zhang et al., 2020). This could account for the reduced flux through the primary metabolic pathways.

The overall observed trends are consistent with the previous reports, which have demonstrated a higher label incorporation during the initial phases of photosynthesis, followed by a slowdown in label incorporation during the later hours (Sharkey et al., 2020; Xu et al., 2022). The kinetic labelling profile across both light regimes points towards a distinct carbon allocation pattern during the mid and late light period (8 and 16 h, respectively) and was thus selected for the detailed mechanistic interpretations. Due to the slower turnover of the TCA cycle (Calvin et al., 1952; Ma et al., 2014; Nogués et al., 2004; Szecowka et al., 2013; Xu et al., 2024) compared to the Calvin-Benson cycle (CBC), the labelling in TCA cycle intermediates is lower under both visible and UV-B light conditions. The overall enrichment pattern across all the measured soluble metabolites points towards an effective carbon assimilation and metabolic flux under visible light, whereas under UV-B, there is a trend of reduced label incorporation.

### Light-phase carbon dynamics highlight mid-light period carbon fixation and late-light period dilution

Due to its robust nature, the mass isotopomer distribution (MID) becomes a valuable component in stable isotope labelling studies (Meiser & Frezza, 2024). As the m+0 isotopomer defines the unlabelled pool of the metabolite mass isotopomers, any decrease in the proportional MID of m+0 over the time points would mean an increase in the ^13^C label incorporation, which would further indicate active involvement of the metabolite in photosynthetic carbon assimilation (Arrivault et al., 2017). The proportion of m+0 isotopomer of each metabolite analysed under visible light conditions provided deeper metabolic insights into the rapeseed seedlings (**Figure 2)**. There is an evident decrease in the proportional MID of the m+0 isotopomer at 8 hours following ^13^CO_2_ exposure under visible light, with statistically significant changes observed in sucrose, serine, shikimic acid, phenylalanine, leucine, isoleucine, valine, ketoglutaric acid, succinic acid, malic acid, glutamic acid, oxoproline and GABA, pointing towards active ^13^C assimilation into these metabolites. This suggests that initially, the assimilated ^13^CO_2_ is incorporated into the sucrose backbone through the CBC via the triose and hexose phosphate pool intermediates (Szecowka et al., 2013). Then, the fixed carbon is channelled to serine, shikimic acid, phenylalanine, leucine, isoleucine, valine and glutamate-GABA metabolism via the central metabolic pathways of glycolysis, PPP (pentose phosphate pathway) and/or the TCA cycle. Upon prolonged exposure to ^13^CO_2_ for 16 hours, the m+0 isotopomer showed a statistically significant increase in the proportional MID in sucrose, serine, isoleucine, ketoglutaric acid, succinic acid, and glutamic acid compared to 8h, pointing towards dilution of the labelled pool, which could be due to the buffering effect or remobilisation of carbon from the unlabelled pools (Koley et al., 2024). At 16h, the m+0 pools of other metabolites, including amino acids (phenylalanine, leucine, valine, threonine and GABA) and shikimate, remained nearly unchanged relative to the 8-hour time point. Valine, citrate and malate were among the three metabolites where we observed a steady decrease in the m+0 isotopomer at both the 8- and 16-hour time points. This can be attributed to the large vacuolar pools having a slower turnover rate (Abadie et al., 2024; Lips & Beevers, 1966a, 1966b; Steer & Beevers, 1967; Szecowka et al., 2013).

**Figure 2:**
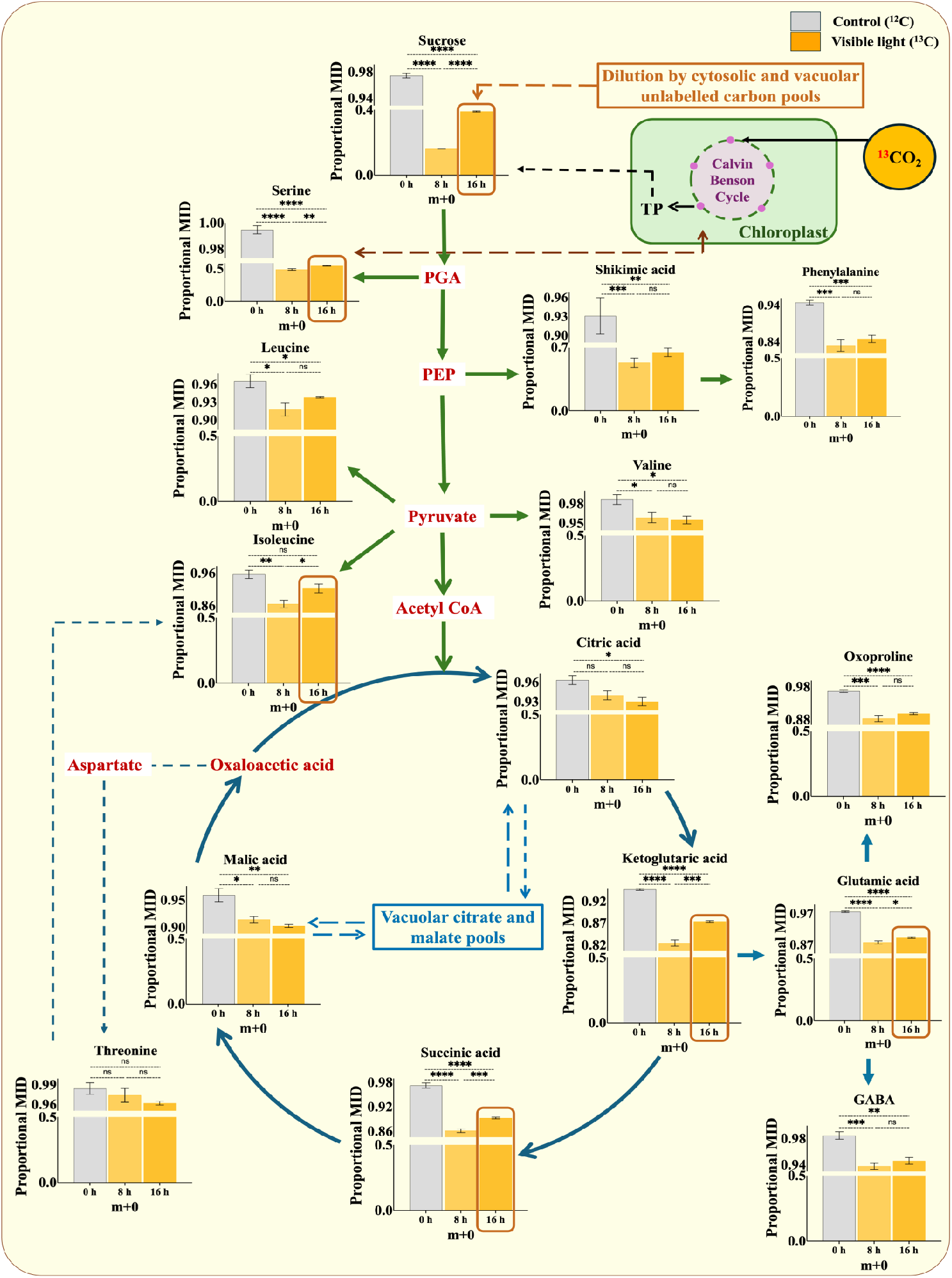
Visible light exhibits progressive enrichment in label incorporation followed by dilution. The proportional mass isotopomer distribution (MID) of the m+0 isotopomer of various metabolites of the central metabolism under visible light at 0h, 8h and 16h is presented. The grey bar represents the starting point (control) before exposing the seedlings to visible light and ^13^CO_2_. The orange bars represent the proportional MIDs of the m+0 isotopomer of metabolites at 8 hours and 16 hours post-^13^CO_2_ exposure under visible light. Pathways such as glycolysis, the TCA cycle, amino acid biosynthesis, GS-GOGAT, and the shikimate pathway are indicated. Error bars are expressed as the standard error of the mean (SEM), where n=4. To determine statistical significance, an unpaired t-test was performed with the P-value adjusted to a threshold of α = 0.05. Significant differences are indicated with an asterisk (*) where p values are reported to 4 decimal places or as non-significant when appropriate. **** - p < 0.0001, *** - p < 0.001, ** - p <0.01, *p < 0.05 and p>0.05 = ns (non-significant).

Collectively, our results demonstrate that there is a dynamic redistribution of the newly fixed ^13^CO_2_ through central metabolism under visible light during the mid- and late-light periods. While the majority of the central metabolites are sustained by active ^13^CO_2_ fixation during the mid-light period (8h), most of these metabolites exhibit an active remobilisation of carbon from the pre-existing unlabelled pools by 16h, except valine, citrate and malate, where, due to the large size of the unlabelled vacuolar pools, the label incorporation seems to be minimal but constantly increasing due to an even slower turnover.

### Light-to-dark metabolic transition in *B. napus* seedlings exhibited remobilisation of photosynthetically fixed ^13^C, supporting in vivo synthesis of shikimic acid and intermediates of the TCA cycle and glutamate-GABA metabolism in the dark

The photosynthetically assimilated ^13^CO_2_ during the light phase was also found to contribute to the increase in the average ^13^C levels in the metabolites during the dark phase. Specifically, the labelling in the metabolites at 24 hours (dark period) was higher in seedlings subjected to dark conditions after receiving ^13^CO_2_ for 16 hours under visible light (**Figure 3)**. During the dark period (night), plants do not photosynthesise, i.e., photosynthetic carbon assimilation stops, and carbon supply to the cytosol occurs mainly in the form of hexose phosphates that are derived from sucrose or starch metabolism, which have complementary yet contrasting functions (Dohler, 1989; Graf & Smith, 2011; Szecowka et al., 2013; Sulpice et al., 2014; Scialdone & Howard, 2015; MacNeill et al., 2017; Wacker et al., 2025).

**Figure 3:**
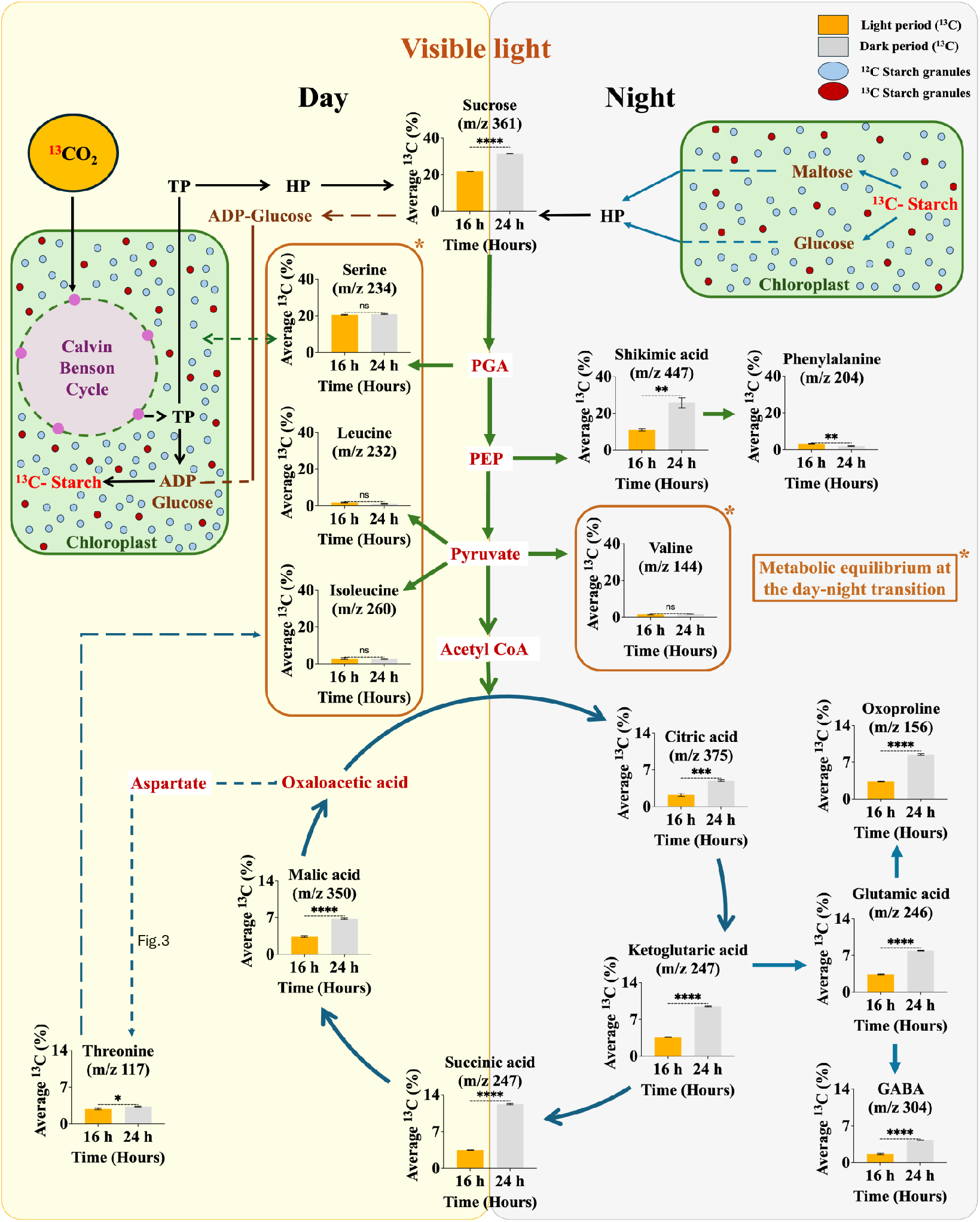
^13^CO_2_ label incorporation into the storage photosynthate and further utilisation of the labelled pool during the light-to-dark transition under visible light conditions. The average 13C incorporation (%) is plotted against the day-night transition (16h and 24h), denoted by orange and grey bars, respectively. Error bars are represented by the standard error of the mean, where n=4. To determine statistical significance, unpaired t-tests were performed, with the P-value adjusted to a threshold of α = 0.05. Significant differences are indicated with an asterisk (*) where p values are reported to 4 decimal places or as non-significant when appropriate. **** - p < 0.0001, *** - p < 0.001, ** - p <0.01, *p < 0.05 and p>0.05 = ns (non-significant). TP = Triose Phosphates and HP = Hexose Phosphates.

It is known that in plants, starch produced during the day is broken down at night into its constituent sugars and sugar alcohols (Koley et al., 2024). The breakdown of starch in the plastid results in a build-up of sucrose in the cytosol. In this manner, respiration at night favours sucrose biosynthesis even in the absence of photosynthesis by utilising the carbon that was stored as starch during the CBC through proper carbon partitioning. This explains the increased levels of ^13^C labels in sucrose under visible light conditions at the 24-hour time point (dark period) (**Figure 3)**. The availability of high ^13^C sucrose reserves fuels all downstream metabolic processes, resulting in higher levels of average ^13^C labels in the glycolytic, TCA, and glutamate-GABA cycle intermediates involving serine, shikimic acid, citric acid, ketoglutaric acid, succinic acid, malic acid, threonine, glutamic acid, GABA and oxoproline supporting the in vivo biosynthesis of these metabolites.

The amino acid pool (leucine, isoleucine, and valine) was maintained at constant levels of ^13^C label at both 16- and 24-hour time points due to recycling with the older ^12^C pools, highlighting an equilibrium in their metabolism in both day and night conditions (Abadie et al., 2024). Also, serine showed no significant changes in the pattern of label incorporation during the light-dark transition, potentially due to an isotopic equilibrium with the relatively large intracellular unlabelled serine pools (Sharkey et al., 2020). Phenylalanine showed a statistically significant decrease in the label incorporation in the dark, which could be due to a reduced carbon flow via the shikimic acid node, its increased utilisation during the dark or due to dilution by the unlabelled precursors (Abadie et al., 2024). Meanwhile, threonine showed a slight increase in the average ^13^C incorporation at 24h.

The label incorporation in the TCA cycle and glutamate metabolism intermediates also displayed a trend of higher label incorporation during the dark. Labelling in citrate and glutamate increased during the dark period (at 24 h) compared to the light period (at 16 h). Typically, it is observed that the average ^13^C in citrate and glutamate during the light phase was comparatively less than that of sucrose and shikimate, likely due to the contributions of pre-existing metabolic pools (Abadie & Tcherkez, 2019; Lawlor & Fock, 1977). The increased labels in citrate and glutamate during the dark indicate their de novo synthesis from the stored assimilated metabolite precursors formed via ^13^CO_2_ fixation (Abadie et al., 2024; Tcherkez et al., 2008, 2012). Further, oxoproline and GABA exhibited a decent label incorporation that increased from 3.32% and 1.6% at 16h to 8.48% and 4.27% at 24h. These observations indicate that the TCA cycle and glutamate-associated pathways of GABA and oxoproline operate smoothly with the breakdown and metabolism of stored ^13^C reserves during the night. Based on these observations, it would be interesting to explore the metabolic reprogramming in *B. napus* seedlings at the light-dark transition under UV-B.

### Delayed ^13^CO_2_ assimilation and conserved carbon pools characterise UV-B-induced metabolic adjustments

On exposure to UV-B for 16 h, *Brassica napus* seedlings exhibited decreased ^13^CO_2_ assimilation and altered metabolism in comparison to visible light exposure **(Figure 2 & 4)**. The proportions of unlabelled metabolites are represented by m+0, and their dilution levels can be used as a readout to correlate with the ^13^C incorporation in other mass isotopomers. It is evident from the extent of the unlabelled metabolite pools that carbon fixation is suppressed, and there is a pattern of delayed label incorporation under UV-B light **(Figure 4)**. Although there is reduced ^13^C incorporation, the channelisation of assimilated carbon follows a trend different from the one observed in visible light **(Figure 2)**. At 8h, there is a significant decrease in the proportional MID of the m+0 isotopomer in sucrose, serine, shikimic acid, TCA cycle intermediates (citric acid, ketoglutaric acid and succinic acid) and the intermediates of the glutamate-GABA metabolism (glutamic acid, GABA and oxoproline). This observation suggests that, despite the metabolic slowdown, ^13^CO_2_ fixation persists under UV-B, and the assimilated carbon is being channelled via the CBC and TCA cycle towards the nitrogen assimilation routes.

**Figure 4:**
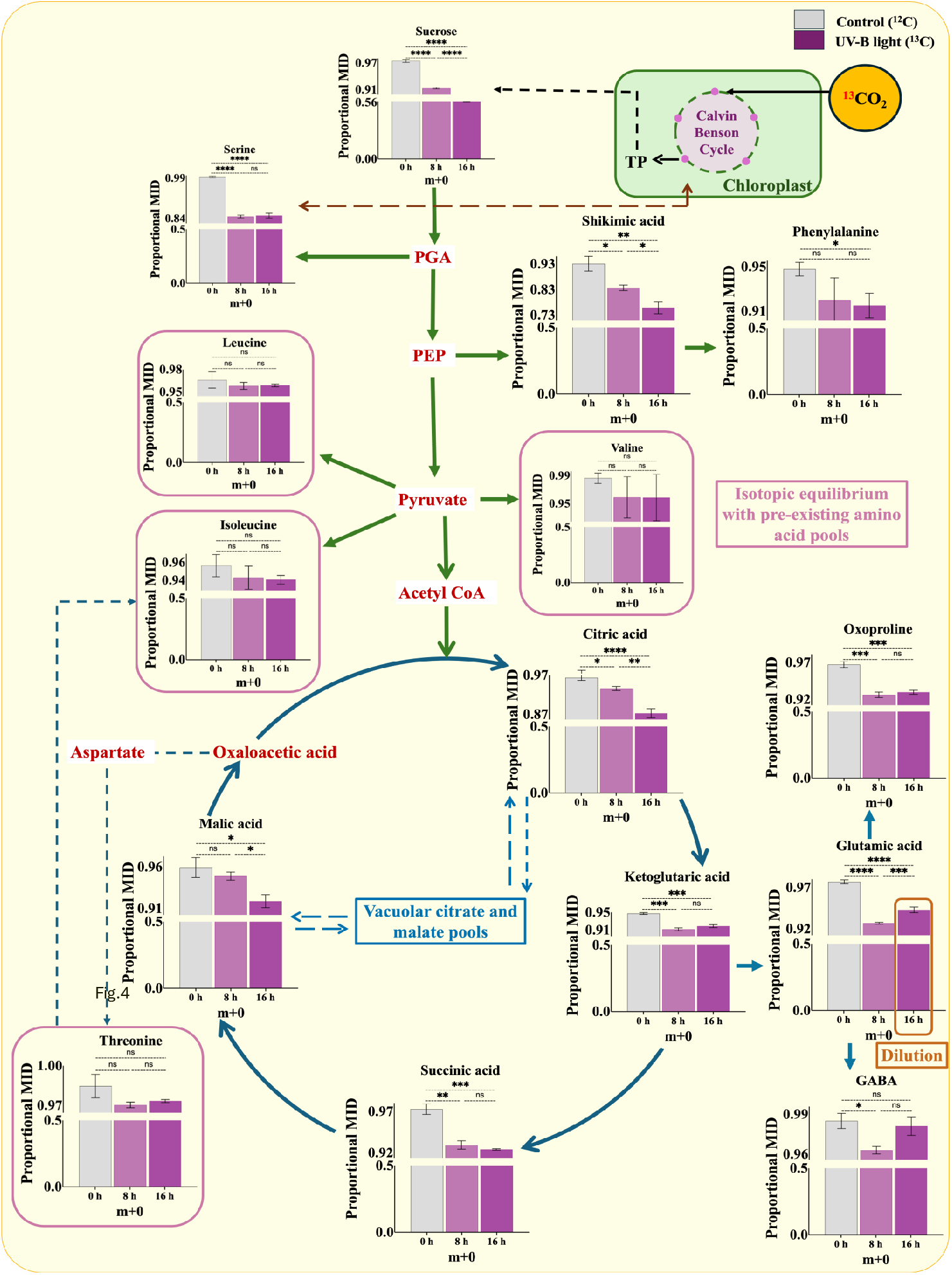
A conservative carbon flux is maintained under UV-B. The proportional mass isotopomer distribution (MID) of the m+0 isotopomer of various metabolites of the central metabolism under visible light at 0h, 8h and 16h is presented. The grey bar represents the starting point (control) before exposing the seedlings to UV-B light and ^13^CO_2_. The violet bars represent the proportional MIDs of the m+0 isotopomer of metabolites at 8 hours and 16 hours post-^13^CO_2_ exposure under UV-B. Pathways such as glycolysis, the TCA cycle, amino acid biosynthesis, GS-GOGAT, and the shikimate pathway are indicated. Error bars are expressed as the standard error of the mean (SEM), where n=4. To determine statistical significance, an unpaired t-test was performed with the P-value adjusted to a threshold of α = 0.05. Significant differences are indicated with an asterisk (*) where p values are reported to 4 decimal places or as non-significant when appropriate. **** - p < 0.0001, *** - p < 0.001, ** - p <0.01, *p < 0.05 and p>0.05 = ns (non-significant).

During the transition from 8 to 16h, sucrose exhibited a further significant increase in the labelled isotopomer as indicated by the decrease in the proportional MID of the m+0 isotopomer. Similar significant changes are also observed in the m+0 isotopomer of shikimate. In contrast, the proportional MID of the other glycolytic intermediate-derived metabolites remained largely unchanged, indicating utilisation of carbon from pre-existing unlabelled pools (Xu et al., 2022). The incorporation of the label in sucrose showed a steep spike, as observed by the decrease in the proportional MID of the m+0 isotopomer from 0.91 at 8 hours to 0.56 at 16 hours. This response in label incorporation points towards the delayed but strong ^13^CO_2_ assimilation in sucrose. Due to the isotopic equilibrium of the large serine pools, no changes were observed in serine labels during this photoperiod. Due to the recycling with their older ^12^C pools, amino acids such as phenylalanine, leucine, isoleucine, valine, and threonine maintained an equilibrium in their metabolism throughout the light period (Abadie et al., 2024). These observations highlight the delayed and limited de novo biosynthesis of sucrose via ^13^CO_2_ assimilation under UV-B during the mid and late light period. It is evident from the observations that newly fixed ^13^CO_2_ didn’t contribute to the de novo biosynthesis of leucine, isoleucine, valine and threonine. However, limited de novo biosynthesis of phenylalanine occurred towards the late light period.

The TCA intermediates, including citric acid, ketoglutaric acid and succinic acid, showed a decrease in the m+0 isotopomer’s proportional MID at 8h. During the transition from 8 to 16h, citric acid and malic acid showed significant label incorporation. These trends point towards the differential flow and temporal redistribution of assimilated carbon from CBC to the TCA cycle. Glutamic acid, oxoproline and GABA also showed a similar trend of label incorporation at 8h under UV-B exposure. Glutamic acid showed an increase in the m+0 fraction at 16 hours, suggesting replenishment from the unlabelled nitrogen-associated pools (Abadie et al., 2017). These observations suggest that UV-B does not completely inhibit photosynthesis; rather, it restricts the rate of carbon assimilation (Dohler, 1989). Although very little label incorporation occurs in the TCA cycle metabolites and glutamate-GABA metabolic cycle intermediates under UV-B light (**Figure 1)**, the de novo biosynthesis of citric acid, ketoglutaric acid, succinic acid, glutamic acid, oxoproline and GABA is still favoured at the mid light period. Moreover, citric acid and malic acid continue their de novo biosynthesis even in the late light period. Glutamic acid was the only amino acid that showed a statistically significant increase in the m+0 isotopomer at 16h, which points towards its dilution by the pre-existing unlabelled carbon pools. Due to this buffering effect, the m+0 proportions in oxoproline and GABA remained nearly constant at the mid and late light periods. Therefore, the carbon metabolism is affected differently by UV-B light, leading to a shift in the metabolism of the central metabolites (Y. Sun et al., 2022; X. Zhang et al., 2018).

### Dark metabolism relies on the mobilisation of pre-existing ^12^C pools under UV-B; however, limited de novo biosynthesis of ketoglutarate, succinic, malic acid and GABA also occurs

Although the average ^13^CO_2_ incorporation in metabolites for the UV-B-treated samples was lower **(Figure 1)**, the labelling pattern at 24 hours showed a drastic difference that was opposite to the trend observed under visible light. The labelling at 16 hours was higher for the glycolytic intermediates compared to the 24-hour time point (**Figure 5)**. This is because UV-B already reduces the photosynthetic efficiency (Kataria et al., 2014) during the light hours, thereby reducing the carbon storage in the form of labelled storage product (possibly starch). A similar trend was also observed in a study by Dohler (1989), where they reported that UV-B exposure led to a decrease in ^14^CO_2_ assimilation. And even if a small amount of labelled ^13^CO_2_ gets stored in the form of starch, there might be a compartmental blockage in the plastid-cytosol transport of hexose phosphates during UV-B exposure. There might be a stress-induced shift in metabolic priorities under UV-B, which favours repair and ROS detoxification over biosynthesis (Shi & Liu, 2021; Tan et al., 2023; Xue et al., 2022), further limiting the ^13^C incorporation into metabolites. There are certain metabolites like ketoglutaric acid, succinic acid, and GABA, which showed a trend of higher ^13^C label (%) at 24h compared to 16h, pointing towards their de novo biosynthesis. This suggests that UV-B rewires the metabolic activities at different time points, diverting the flux from ideal carbon fixation to stress-induced responses by actively channelling the limited pool of labelled carbon and utilising the pre-existing unlabelled carbon pools. Therefore, an altered source-to-sink dynamics exists in the metabolic phenotype under UV-B stress, pointing towards dependence on pre-existing ^12^C sucrose pools.

**Figure 5:**
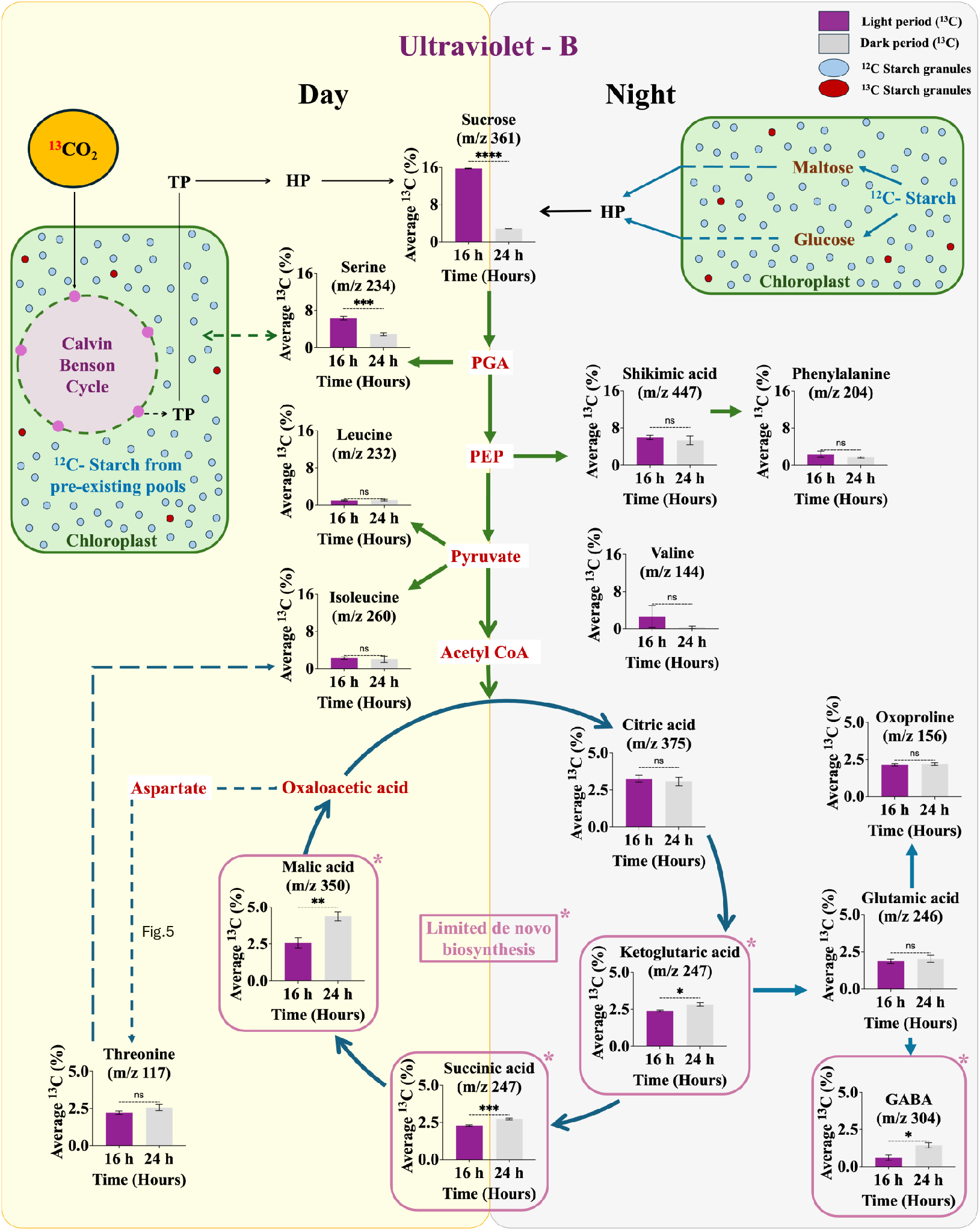
Utilisation of carbon from pre-existing ^12^C-sucrose pools under UV-B during light-to-dark transition. The average ^13^C label incorporation (%) in various metabolite fragments is represented under UV-B light at 16h and 24 h, violet and grey bars. Error bars are represented by the standard error of the mean, where n=4. To determine statistical significance, unpaired t-tests were performed, with the P-value adjusted to a threshold of α = 0.05. Significant differences are indicated with an asterisk (*) where p values are reported to 4 decimal places or as non-significant when appropriate. **** - p < 0.0001, *** - p < 0.001, ** - p <0.01, *p < 0.05 and p>0.05 = ns (non-significant). TP = Triose Phosphates and HP = Hexose Phosphates.

### Integrative isotopomer heatmap reveals reprogrammed central metabolism under visible and UV-B light

**Figure 6.** gives a visual representation of the kinetic light-dark transition. With the decrease in the m+0 isotopomer (unlabelled portion), there is a concurrent increase in the m+1, m+2 and other isotopomers, indicating an active time-dependent label incorporation into central metabolites. There is an active incorporation of the labelled ^13^CO_2_ into most of the metabolites under visible light; however, the rate of labelling is lower under UV-B light.

**Figure 6:**
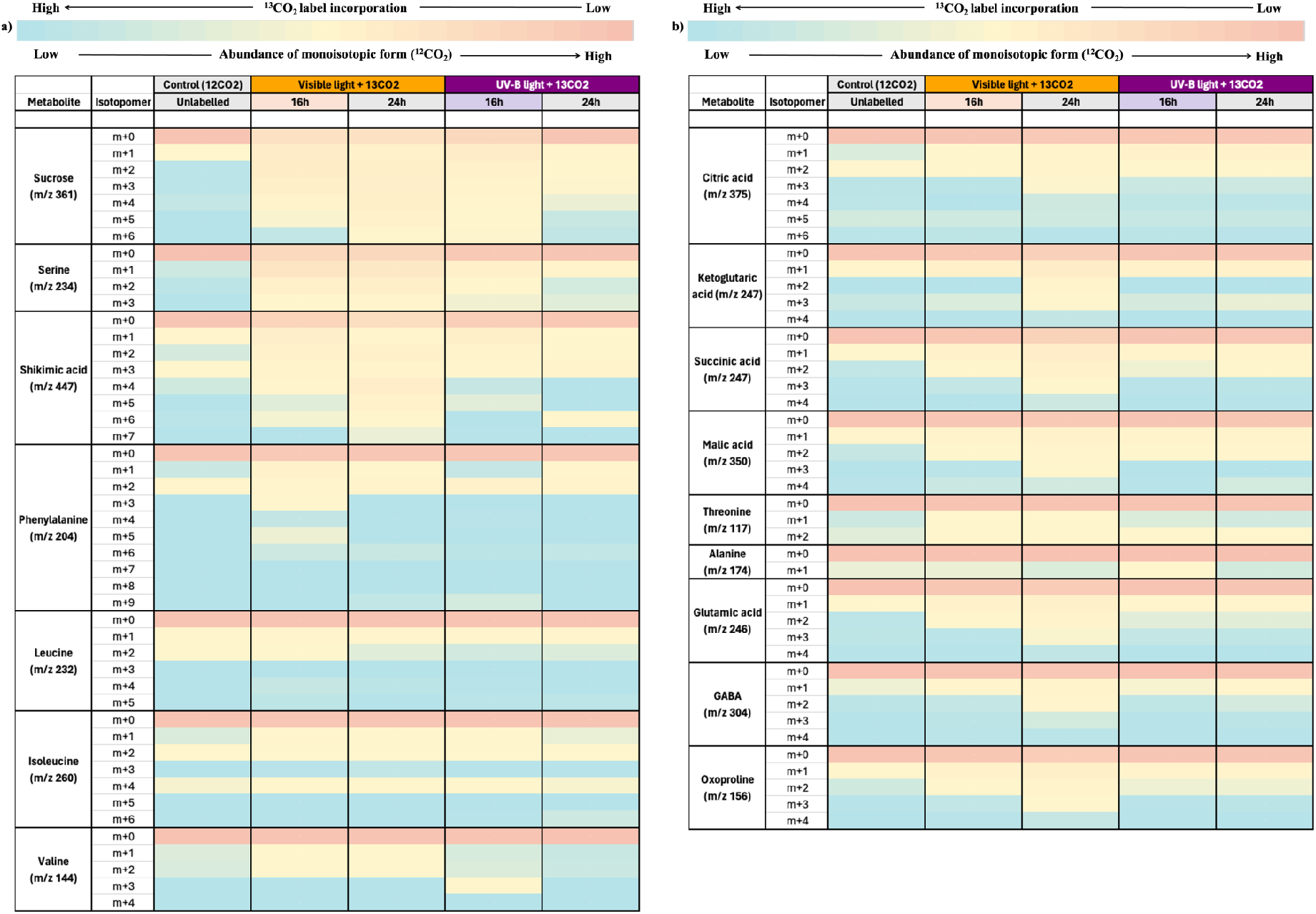
Heatmap illustrating the proportional mass isotopomer distribution of various metabolite fragments following ^13^CO_2_ labelling under visible and UV-B light exposure for 16 and 24h. The metabolite fragments were analysed for label incorporation under visible and UV-B light conditions. The colour gradient indicates the relative proportion of monoisotopic form (^12^C) and the extent to which its dilution occurs during the incorporation of ^13^CO_2_. Blue represents a low abundance of ^12^C and thus a higher ^13^C label incorporation, and red represents high ^12^C abundance and a lower ^13^C label incorporation. A) represents the label incorporation in the CBC derived intermediates, and b) represents the label incorporation in the TCA cycle and glutamate-GABA metabolism intermediates

When subjected to visible light, most of the glycolytic intermediate-derived metabolites, including sucrose, serine and shikimic acid, showed a large drop in m+0. However, metabolites of the TCA cycle (citric acid, ketoglutaric acid, succinic acid, malic acid) and glutamate-GABA metabolism (glutamic acid, GABA, oxoproline) had a moderate decrease in the m+0 fraction. In contrast, the amino acids maintained a nearly constant pool of the unlabelled metabolites. This suggests that there is an active photosynthetic assimilation into the metabolites of glycolysis, the TCA cycle, and the glutamate-GABA metabolism intermediates. The amino acid pool, however, remains in constant sync with the pre-existing unlabelled pools. The decrease in the m+0 fraction was higher in the dark (24h) compared to the light period (16h), as observed by the increase in the fraction of other isotopomers. This suggests that carbon assimilation and storage during the day (16h) and redistribution at night (24h) are supported under visible light.

Under UV-B light, the label incorporation patterns are altered. We observed reduced and delayed label incorporation into the metabolites, resulting in a constant proportion of the m+0 isotopomer across the majority of the metabolites. Sucrose, serine and shikimic acid showed a moderate decrease in the unlabelled fraction at 16h, but during the dark period (24h), the m+0 isotopomer exhibited a further increase, suggesting that the fixed ^13^CO_2_ is getting consumed during the day hours only, and the dark metabolism is mostly supported by pre-existing unlabelled pools.

## Conclusion

The photosynthetic carbon fixation and allocation of photosynthates through central metabolism are driven by light, and therefore, the quality of light plays a decisive role in shaping the metabolic phenotype. In the present study on rapeseed seedlings, we explored a wide range of metabolites under visible and UV-B light using metabolic profiling and ^13^CO_2_ mapping. Metabolism is an inherently dynamic and complex process, and metabolomics provides only a static snapshot of this process (Meiser & Frezza, 2024). Therefore, we employed kinetic ^13^CO_2_ mapping to capture a dynamic view of metabolism under visible and UV-B light by connecting light-to-dark transitions with carbon partitioning strategies. There was an observable difference in resource partitioning under both light conditions. Kinetic ^13^CO_2_ mapping revealed that the light condition (visible and UV-B) induced divergent metabolic phenotypes. Under visible light, the seedlings maintained an active basal metabolic flux indicative of active carbon assimilation, where they utilised the assimilated ^13^CO_2_ towards normal growth and storage activities. In contrast, when exposed to UV-B, there was a change in the physiological response from growth towards protection. This reorientation suggests metabolic flexibility under different light conditions.

Our findings highlight that kinetic ^13^CO_2_ mapping captures insights beyond steady-state metabolomics. Visible light supported the dynamic redistribution and storage of labelled carbon during the mid to late light hours, while the metabolism in the dark favoured the breakdown of labelled storage product into sucrose. The de novo biosynthesis of shikimic acid, TCA cycle intermediates (citric acid, ketoglutaric acid, succinic acid and malic acid), and the intermediates of glutamate-GABA metabolism (glutamic acid, oxoproline and GABA) were also largely supported in dark metabolism. Amino acid pools of serine, leucine, isoleucine and valine, however, remained in a state of metabolic equilibrium with their unlabelled pools during the day-night transition photoperiod.

The majority of rapeseed metabolism was sustained by recycled or pre-existing pools under UV-B light; however, certain TCA cycle intermediates, including keto glutarate, succinic acid and malic acid, along with GABA, exhibited limited de novo biosynthesis in the dark. Consistent with previous reports, which highlight the role of TCA cycle intermediates in energy metabolism (Lingwan et al., 2024), we observed a trend of lower label incorporation into TCA cycle intermediates under UV-B. This highlights the reorientation in metabolism wherein the plant restricts its carbon flow away from energy production towards stress-responsive metabolism by prioritising stress response over growth. The delayed labelling patterns and conserved carbon pools (indicated by the proportional MID of the m+0 isotopomer) highlighted that the seedling engaged in slightly downregulated but active photosynthetic activity during the light period, while metabolism in the dark relied on stored unlabelled or recycled carbon sources predominantly.

## Supplementary data

Additional information included here

**Supplementary Table S1:** Average ^13^C incorporation in all the metabolite fragments generated in this study.

**Supplementary Table S2:** Valid TMS-Fragments selected for ^13^CO_2_ mapping in rapeseed seedlings.

**Supplementary Table S3:** GC-MS characteristics of soluble metabolites detected in *Brassica napus* var. *sheetal* seedlings.

**Supplementary Table S4:** The Mass Isotopomer Distributions (MIDs) of all valid metabolite fragments generated in this study.

**Supplementary Figure S1:** Labelling pattern in the central metabolic pathway intermediates showing the average ^13^C incorporation (%) in TMS fragments. Error bars are represented by the standard error of the mean, where n=4. a) Indicates the valid fragments with kinetic ^13^C enrichment levels exceeding the average natural abundance of ^13^C over different time points, and b) indicates the fragments that showed label incorporation but were not included during further analysis

**Supplementary Figure S2:** Overall variation in labelling during the day-night transition.

**Supplementary Figure S3:** GC-MS spectra showing the total ion chromatogram (TIC scan) of 8-day-old rapeseed seedlings subjected to visible and UV-B light at 16 and 24-hour time points.

## Acknowledgements

SKM acknowledges the Science and Engineering Research Board (SERB) Early Career Research funding. YT acknowledges the University Grants Commission (UGC) for her PhD fellowship. ML and YP acknowledge the Indian Institute of Technology, Mandi, for their PhD fellowships.

## Author contributions

SKM and ML conceptualised the study. ML and YP performed the experiments. YT analysed the data, made all the figures, tables and prepared the original draft. SKM reviewed and edited the manuscript. SKM also acquired the funding.

## Data Availability

All supporting data are available in the manuscript figures and supplementary materials.

