## Supplementary Figures for "Kinetic ^13^CO_2_ mapping revealed distinct light-dark metabolic transition phenotypes in *Brassica napus* seedlings under visible and UV-B light"

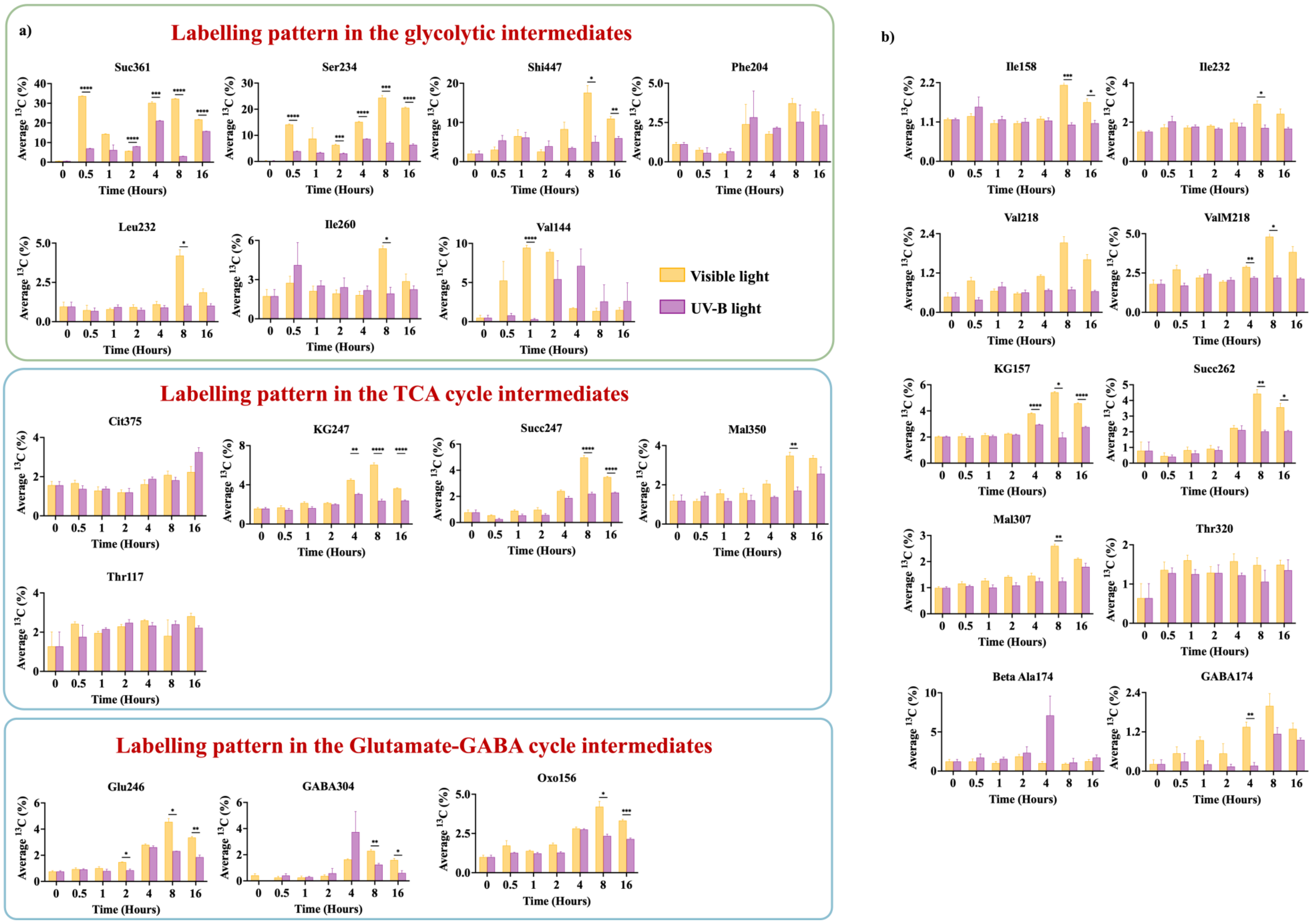


**Supplementary Figure S1-** Labelling pattern in the central metabolic pathway intermediates showing the average 13C incorporation (%) in TMS fragments. Error bars are represented by the standard error of the mean, where n=4. a) Indicates the valid fragments with kinetic 13C enrichment levels exceeding the average natural abundance of 13C over different time points. b) Indicates the fragments that showed label incorporation but were not included during further analysis. To determine statistical significance, unpaired t-tests were performed for multiple comparisons, where the P value was adjusted to a threshold of α = 0.05. Significant differences are indicated with an asterisk (*) where p values are reported to 4 decimal places or as non-significant when appropriate. **** - $p < 0.0001$, *** - $p < 0.001$, ** - $p < 0.01$, *$p < 0.05$ and $p > 0.05$ = ns (non-significant).

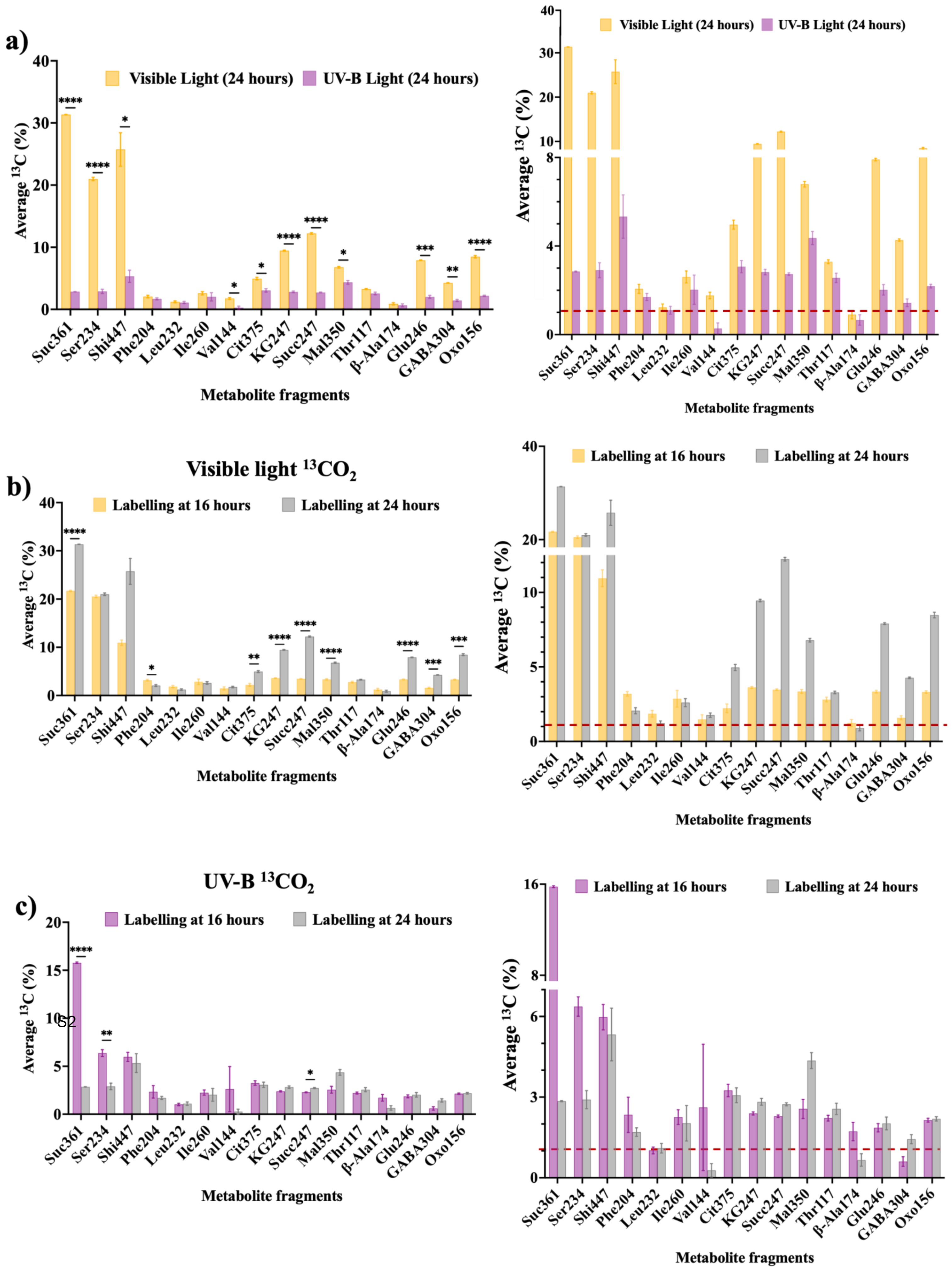


**Supplementary Figure S2: Overall variation in labelling during the day-night transition**. a) represents higher labelling in the metabolites of samples exposed to a dark period after 16 hours of visible light exposure than in samples exposed to a dark period after 16 hours of UV-B exposure. b) represents visible light 16-hour vs 24-hour samples, where label incorporation was higher at 24 hours. c) represents UV-B 16-hour vs 24-hour samples where a higher $^{13}CO_2$ label has been incorporated in the light hours (16 hours), a trend contrary to the visible light condition. Error bars are represented by the standard error of the mean, where n=4. To determine statistical significance, unpaired t-tests were performed for multiple comparisons, with the P-value adjusted to a threshold of $\alpha = 0.05$. Significant differences are indicated with an asterisk (*) where p values are reported to 4 decimal places or as non-significant when appropriate. **** - $p < 0.0001$, *** - $p < 0.001$, ** - $p < 0.01$, *$p < 0.05$ and $p > 0.05$ = ns (non-significant).

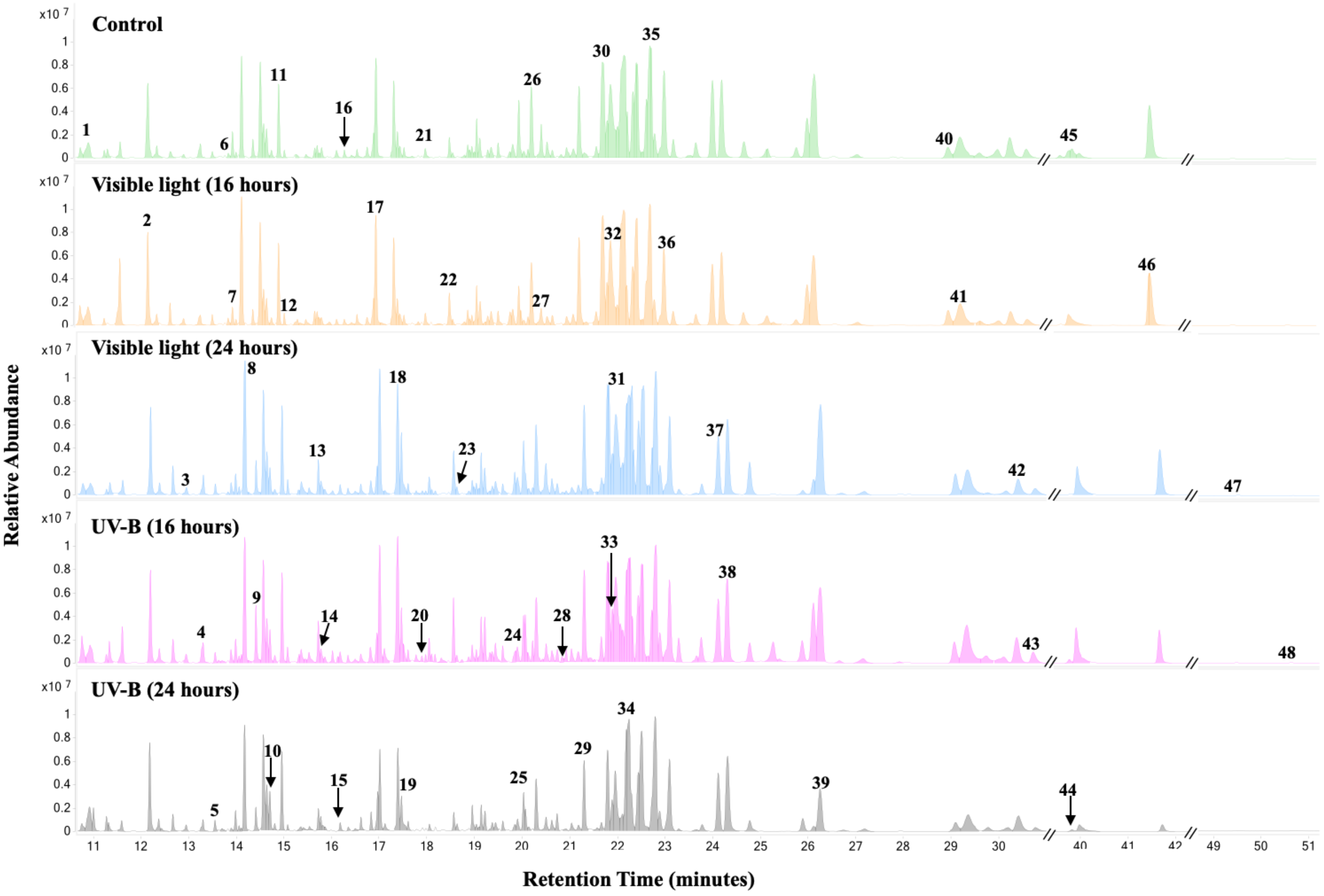


**Supplementary Figure S3: GC-MS spectra showing the total ion chromatogram (TIC scan) of 8-day-old rapeseed seedlings subjected to visible and UV-B light at 16 and 24-hour time points.** The relative peak abundance is plotted against the retention time (minutes). The major metabolites are numbered (Ribitol is used as an internal standard).
